# PATROL1-mediated H^+^-ATPase translocation boosts plant growth under drought by optimizing root and leaf functions

**DOI:** 10.1101/2024.11.04.621996

**Authors:** Naoya Katsuhama, Kazuma Sakoda, Haruki Kimura, Yutaro Shimizu, Yuuki Sakai, Kenji Nagata, Mitsutomo Abe, Ichiro Terashima, Wataru Yamori

## Abstract

Optimizing leaf photosynthesis and root water and mineral uptake in crops during drought is crucial for enhancing agricultural productivity under climate change. Although plasma membrane H^+^-ATPase plays a key role in plant physiological processes, its overexpression alone does not consistently improve growth. While PROTON ATPASE TRANSLOCATION CONTROL 1 (PATROL1) regulates H^+^-ATPase translocation in response to various environmental stimuli in leaves, its function in roots remains largely unknown. Here we show that H^+^-ATPase was coimmunoprecipitated with PATROL1 in roots of *Arabidopsis thaliana*. Micrografting between wild type and *PATROL1* knockout or overexpression lines showed that PATROL1 are indispensable in both shoots and roots, indicating that root uptake and leaf photosynthesis are simultaneous limiting factors for plant growth under drought conditions. *PATROL1* overexpression in whole plants resulted in a 41% increase above wild type in shoot dry weight under drought conditions. These findings highlight the potential of H^+^-ATPase regulation in roots as a new strategy to improve plant productivity, particularly under drought conditions.

## Introduction

Drought presents a major challenge to agricultural production, especially in meeting the food demands of the world’s growing population under climate change ^1^. Yield reduction due to drought occurs when the uptake of water and nutrients, which are essential for photosynthetic assimilation, is restricted ^2,3^. Drought tolerance can take various forms; however, from an agricultural perspective, optimizing water use and capture within the limitation of available soil moisture during gradual and moderate soil water deficits is the most effective adaptation ^4,5^. In this scenario, robust leaf growth and photosynthesis, supported by efficient water and nutrient uptake through enhanced root system architecture and activity, are key traits to minimizing yield penalties or failure ^2,3^. In addition, coregulating photosynthesis and nutrient uptake is advocated as a breeding strategy for improving yield and resource use efficiency even under sufficient irrigation ^6–8^. However, maintaining an excessive root system and its functions consume carbon and energy ^4,9^. Therefore, uncovering root phenotypic plasticity in response to fluctuating soil moisture and nutrient conditions is essential for designing an ideal root system ^10,11^.

Since plasma membrane (PM) H^+^-ATPase plays key roles in plant growth, such as hypocotyl and root elongation, mineral nutrient acquisition, stomatal opening, and environmental stress responses ^12^, modulating PM H^+^-ATPase is a potential strategy to improve root uptake and photosynthesis under drought and resource scarcity ^13^. For instance, overexpression of *Arabidopsis thaliana* H^+^-ATPase 2 (AHA2) in the guard cell promoted light-induced stomatal opening and enhanced growth in *A. thaliana* ^14^ and a *Populus* sp. hybrid ^15^. Moreover, overexpression of *Oryza sativa* H^+^-ATPase 1 in whole plants increased grain yield by simultaneously improving nutrient uptake, photosynthesis, sugar translocation, and nutrient enrichment ^7^. On the other hand, excessive or constitutive PM H^+^-ATPase activity can have adverse effects, including growth impairment, male sterility, necrotic lesions, and severe dehydration, because PM H^+^-ATPase is an ATP-consuming protein with numerous physiological roles ^16,17^. Therefore, both the quantity and timing of PM H^+^-ATPase activity should be precisely regulated to develop drought-tolerant plants while managing associated trade-offs.

Most previous studies of PM H^+^-ATPase regulation focused on transcriptional and post-translational modifications, particularly the phosphorylation and dephosphorylation of autoinhibitory C-terminus residues ^12,13^. PROTON ATPASE TRANSLOCATION CONTROL 1 (PATROL1) was discovered as a unique regulatory factor for *A. thaliana* H^+^-ATPase 1 (AHA1) ^18^. PATROL1 is a plant ortholog of mammalian uncoordinated 13s (Munc13s), and it contains a MUN domain (Supplementary Fig. S1) which is responsible for synaptic vesicle priming in neuronal exocytosis in animals ^18,19^. PATROL1 rapidly tethers an appropriate amount of AHA1 to the plasma membrane of stomatal guard cells in response to changes in light, CO_2_ concentration ([CO_2_]), abscisic acid (ABA), osmotic pressure, and leaf detachment ^20^. Systemic overexpression of *PATROL1* enhances photosynthesis and biomass production under fluctuating light conditions, while maintaining water use efficiency by ensuring rapid stomatal movement ^21^. A recent study using a *patrol1* loss-of-function mutants showed that PATROL1 is also involved in primary root elongation by regulating the root meristematic size ^22^. In addition, shoot growth of wild type plants was reduced when shoot was grafted onto *patrol1* rootstock ^22^ Although PATROL1 is expressed in both shoots and roots ^18,22^, its specific function in roots and their contributions to shoot growth under drought remain unknown.

Here, we examined the distribution pattern and interaction of PATROL1 with AHAs in the roots of *A. thaliana*. We then analyzed the expression and intracellular dynamics of PATROL1, and the root system architecture in wild type, *PATROL1* knockout, and overexpression lines under PEG-simulated drought conditions. Finally, we investigated the distinct roles of PATROL1 in roots and shoots, focusing on nitrogen uptake and photosynthesis under moderate drought stress, aiming at developing a novel strategy for enhancing crop drought tolerance by regulating PM H^+^-ATPase activity.

## Results

### PATROL1 distribution pattern and co-immunoprecipitation with plasma membrane H^+^-ATPase in roots

Overexpression of *PATROL1* (*PATROL1*-OX) promotes photosynthesis induction and growth under fluctuating light conditions owing to rapid stomatal movement ^21^. To investigate other functions of PATROL1, we grew *A. thaliana* wild type (WT), *PATROL1* knockout (*patrol1*), and *PATROL1*-OX plants under elevated [CO_2_], at which stomatal conductance does not limit photosynthesis. Shoot growth of *patrol1* was significantly reduced by 28% relative to WT and by 32% relative to *PATROL1*-OX (Fig. 1a). Although the photosynthetic capacity of *patrol1* increased as [CO_2_] elevated, it remained significantly lower than that of WT and *PATROL1*-OX (Supplementary Fig. S2a right). These results suggests that stomatal limitation is not the sole cause of the impaired CO_2_ assimilation and growth in *patrol1* (Fig. 1a and Supplementary Fig. S2b).

**Fig. 1.**
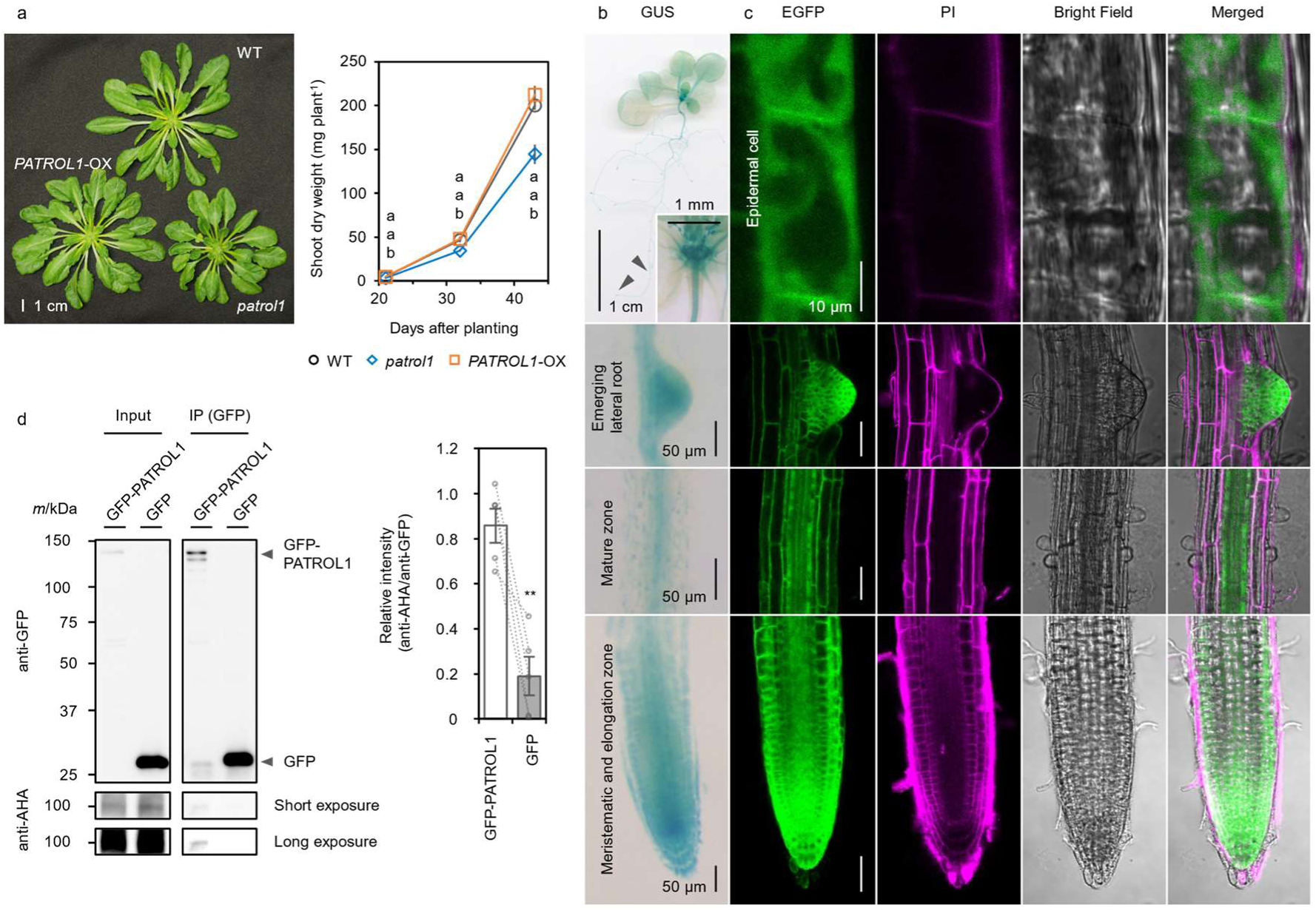
Localization of PATROL1 and interaction with AHAs in the roots of *Arabidopsis thaliana*. **a** Phenotype and shoot dry weight of WT, *patrol1*, and *PATROL1*-OX plants grown at elevated [CO_2_] (1500 μmol mol^−1^) for 43 days. Error bars represent SEM (*n* = 7). Means not sharing any letters are significantly different between genotypes at *p* < 0.05 by Tukey–Kramer test. **b** GUS staining of 2-week-old *gPATROL1:GUS-PATROL1* plants. **c** Subcellular localization of EGFP-PATROL1 in 2-week-old *A. thaliana* primary root. Green, EGFP-PATROL1 localization; magenta cell wall propidium iodide (PI) staining. **d** Association of GFP-PATROL1 and AHAs observed by co-immunoprecipitation (co-IP) assay in the root. Total proteins (input) from the transgenic *A. thaliana* roots producing either GFP-PATROL1 or GFP alone (negative control) were immunoprecipitated with anti-GFP beads. The inputs and immunoprecipitates were immunoblotted with anti-AHA and anti-GFP antibodies. Error bars represent SEM (*n* = 5). ** indicate statistically significant differences by Student’s *t*-test at *p* < 0.01.

We further explored PATROL1 functions besides stomatal regulation. We examined the expression pattern of *PATROL1* in β-glucuronidase (GUS) reporter lines. In these plants, PATROL1 fused to GUS at the N-terminus (GUS-PATROL1) was expressed under the control of the *PATROL1* regulatory sequence. GUS activity was ubiquitous in *gPATROL1:GUS-PATROL1; patrol1* (*gPATROL1:GUS-PATROL1*) seedlings, with higher levels in the vascular bundles, meristematic regions of shoot and primary root, and lateral root primordia (Fig. 1b and Supplementary Fig. S3). In the root elongation and maturation zones, it was moderate in epidermis, cortex, and pericycle (Fig. 1b). We also investigated the subcellular localization of the PATROL1 protein, expressed under the control of the *PATROL1* regulatory sequences, by observing enhanced green fluorescent protein (EGFP)-tagged PATROL1 in *gPATROL1:EGFP-PATROL1; patrol1* (*gPATROL1:EGFP-PATROL1*) plants. The expression pattern of EGFP-PATROL1 in roots was very similar to that of GUS-PATROL1, and EGFP fluorescence was detected in almost all cells, including root epidermal cells, in which EGFP-PATROL1 appeared in the cytosol along the plasma membrane (Fig. 1c).

In stomatal guard cells and subsidiary cells, PATROL1 plays key roles in regulating the trafficking of AHAs to and from the plasma membrane ^18,23^. However, it is unclear whether it also interacts with AHAs in the root. To better understand the relationship between PATROL1 and AHAs in underground parts, we performed co-immunoprecipitation (co-IP) experiments using protein extracts from roots of GFP-PATROL1 or GFP plants. Coimmunoprecipitation of substantial AHA2 (and possible other AHAs: Supplementary Table 2) with GFP-PATROL1, but virtually none with the free GFP control (Fig. 1d), suggests that PATROL1 interacts with AHAs in root cells. Taken together, our results support a model in which plant growth could be affected by the interaction of PATROL1 and PM H^+^-ATPase in roots.

### PATROL1 is essential for root elongation and branching under PEG-simulated drought

To test our hypothesis on the role of PATROL1 in optimizing root growth under soil moisture deficit, we examined how transgenic *A. thaliana* roots respond under hyperosmotic stress. Seedlings were grown on 1/2MS medium for 5 days and then transferred to 1/2 MS medium treated with or without 30% (w/v) PEG6000 for another 10 days. In PEG medium, *PATROL1*-OX plants relatively sustained roots and shoots growth, but that of WT and *patrol1* plants was inhibited (Fig. 2a right). In the control medium, growth of *PATROL1*-OX was comparable to that of WT, but that of *patrol1* was somewhat smaller (Fig. 2a left). The components of the root system and the relative growth of PEG treatment compared to the mean of control were analyzed to assess the impact on root growth. In the control medium, root length and number of WT and *PATROL1*-OX were statistically equivalent, but dose of *patrol1* were significantly less (Fig.2b left). Under hyperosmotic stress, they were significantly greater in *PATROL1*-OX than in WT and significantly less in *patrol1* (Fig. 2b right). The primary root length of WT decreased by 12% of the control, lateral root length by 54%, and lateral root number by 31% (Fig. 2c). Interestingly, those of *patrol1* and *PATROL1*-OX decreased by 4% and 4%, 30% and 32%, and 14% and 6%, respectively, of WT (Fig. 2c). The high relative shoot and root growth in *patrol1* and *PATROL1*-OX plants under water deficit suggests that *patrol1* plants showed survival behaviour, while *PATROL1*-OX plants adopted an escape response (Fig. 2c). Chlorophyll fluorescence imaging confirmed that hyperosmotic treatment reduced the quantum yield of PSII (Y(II)) and increased non-photochemical quenching (NPQ) of *patrol1*, while all three lines showed similar trends in the control (Fig. 2a). Overall, *PATROL1*-OX had 53% greater shoot fresh weight and 37% greater root fresh weight than WT, while *patrol1* had 33% and 41%, respectively, less (Fig. 2b right).

**Fig. 2.**
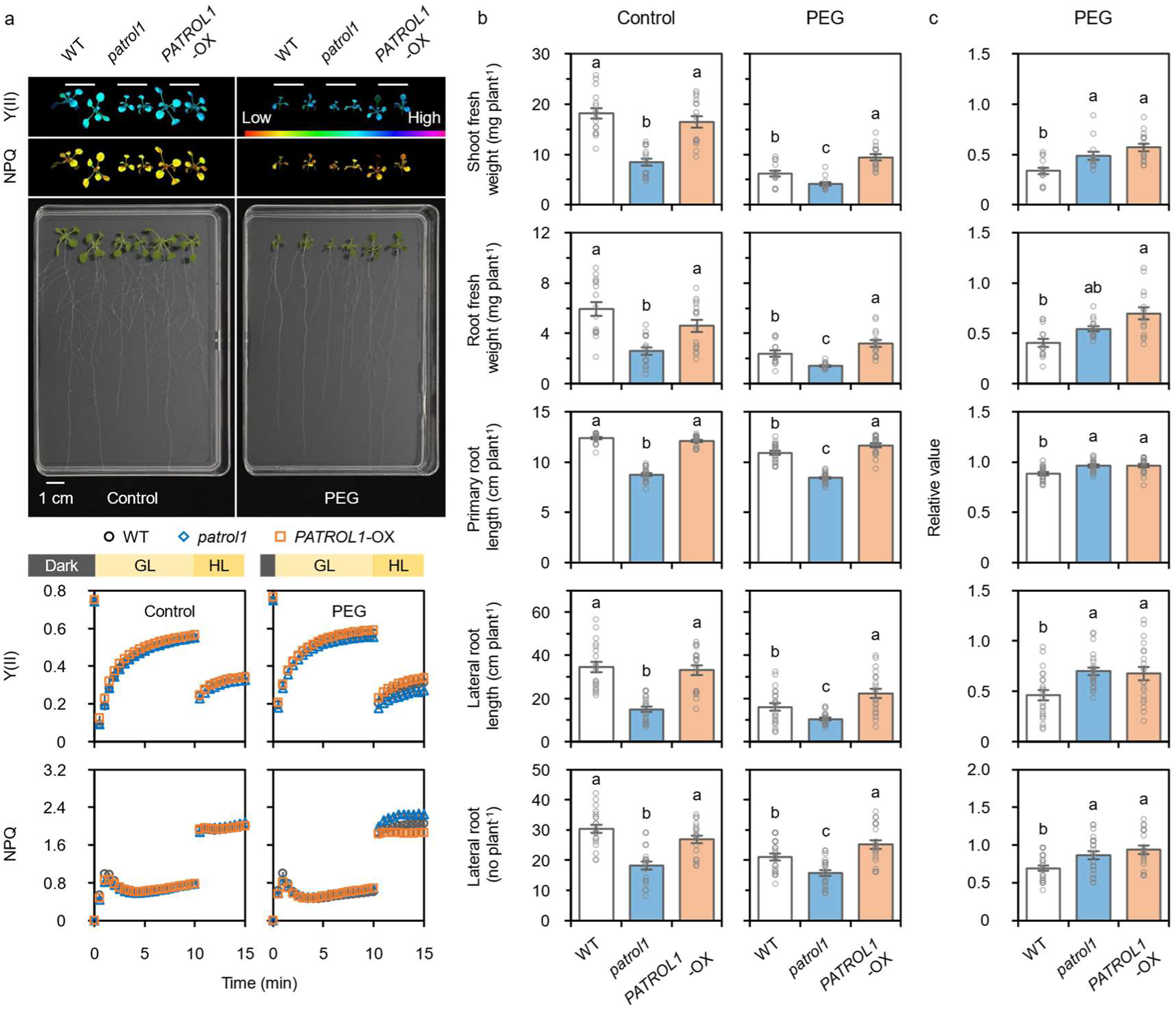
Shoot and root growth of *PATROL1* transgenic plants under PEG-simulated drought conditions. **a** Upper: Phenotype and photochemical activities of WT, *patrol1*, and *PATROL1*-OX plants grown on 1/2 MS medium treated with or without 30% (w/v) PEG 6000 for 10 days. Post-illumination induction of the quantum yield of PSII (Y(II)) and non-photochemical quenching (NPQ), measured under growing light (GL, PPFD of 100 µmol m^−2^ s^−1^) and high light (HL, PPFD of 500 µmol m^−2^ s^−1^) intensities after overnight dark adaptation. Error bars represent SEM (*n* = 8–12). **b** Shoot and root fresh weights, root length and numbers of treated and untreated plants. **c** Values in PEG treatment relative to the average control values were calculated as an indicator of stress tolerance. Error bars represent SEM (*n* = 20–22). Means not sharing any letters are significantly different between genotypes at *p* < 0.05 by Tukey–Kramer test.

We next sought to determine how hyperosmotic stress affects the expression and intracellular dynamics of *PATROL1* in roots. Quantitative reverse-transcription PCR (RT-qPCR) analysis revealed that *PATROL1* expression was slightly upregulated in roots after exposure to PEG (Fig. 3a). Moreover, 30-min exposure to osmotic stress increased EGFP-PATROL1 bodies ^18^ in the peripheral cytoplasm of root epidermal cells within the elongation zone of transgenic plants expressing EGFP-PATROL1 driven by its native promoter (Fig. 3b and Supplementary Fig. S4). This increase in EGFP-PATROL1 bodies suggests that the localization of PATROL1 in roots is regulated in response to hyperosmotic stress, as well as transcriptionally. Notably, the expression levels of five other Domain of Unknown Function 810 (DUF810) and AHA member genes were consistent across WT, *patrol1*, and *PATROL1*-OX plants under control condition (Supplementary Fig. S5).

**Fig. 3.**
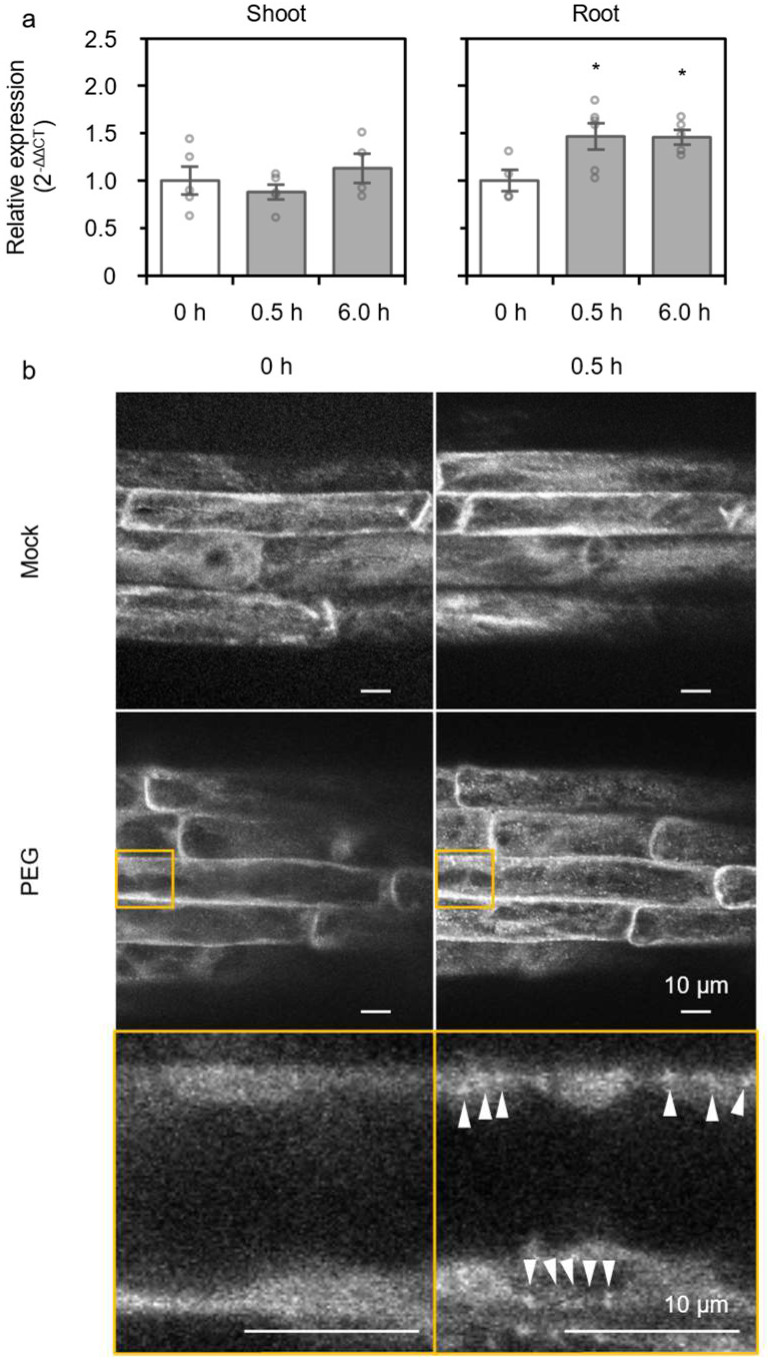
Regulation of root *PATROL1* transcripts and protein localization in response to PEG-induced hyperosmotic stress. **a** Quantitative reverse-transcription PCR analysis of *PATROL1* in shoot and root. 19-day-old *A. thaliana* were treated with 30% (w/v) PEG 6000 for designed times. The expression level of *PATROL1* at 0.0 h of treatment was normalized as 1.0. Error bars represent SEM (*n* = 4–5). * Significant differences between means at *p* < 0.05 by Dunnett’s test. **b** Root epidermal cells from 1-week-old *gPATROL1:EGFP-PATROL1* transgenic plants were treated with liquid 1/2 MS medium (mock) or liquid 1/2 MS medium with 10% (w/v) PEG 8000 for 0.5 h. Arrowheads indicate EGFP-PATROL1 bodies in endosomes responding to osmotic stress. Maximum projection images were generated from five serial Z-slices with a 1-µm interval before and after mock or PEG treatment. Enlarged images of the areas enclosed by yellow squares were taken from a single slice. Bars = 10 mm.

### Overexpression of *PATROL1* enhances root water and nitrogen uptake and improves photosynthesis under moderate drought conditions

To assess the contribution of PATROL1 to plant growth under soil water deficit, we grew WT, *patrol1*, and *PATROL1*-OX plants together in a single pot for 16 days with sufficient water, and then withheld watering for 21 days (Fig. 4a). While the soil volumetric water content (VWC) of the control ranged between 0.50 m^3^ m^-3^ and 0.61 m^3^ m^-3^, those of the treatment pots gradually decreased from 0.60 m^3^ m^-3^ to 0.07 m^3^ m^-3^ due to evapotranspiration (Fig. 4b). Under drought, the shoot dry weight and projected leaf area of *PATROL1*-OX were 41% and 34%, respectively, higher than WT (Fig. 4c right), and those of *patrol1* were 31% and 18% lower than WT (Fig. 4c right). In the control, there were no significant differences in either among the genotypes (Fig. 4c left). In the control, carbon isotope discrimination (δ^13^C), which reflects stomatal closure and CO_2_ diffusion during growth, was identical among genotypes, but under drought, it increased significantly in *patrol1* above WT (Fig. 4d).

**Fig. 4.**
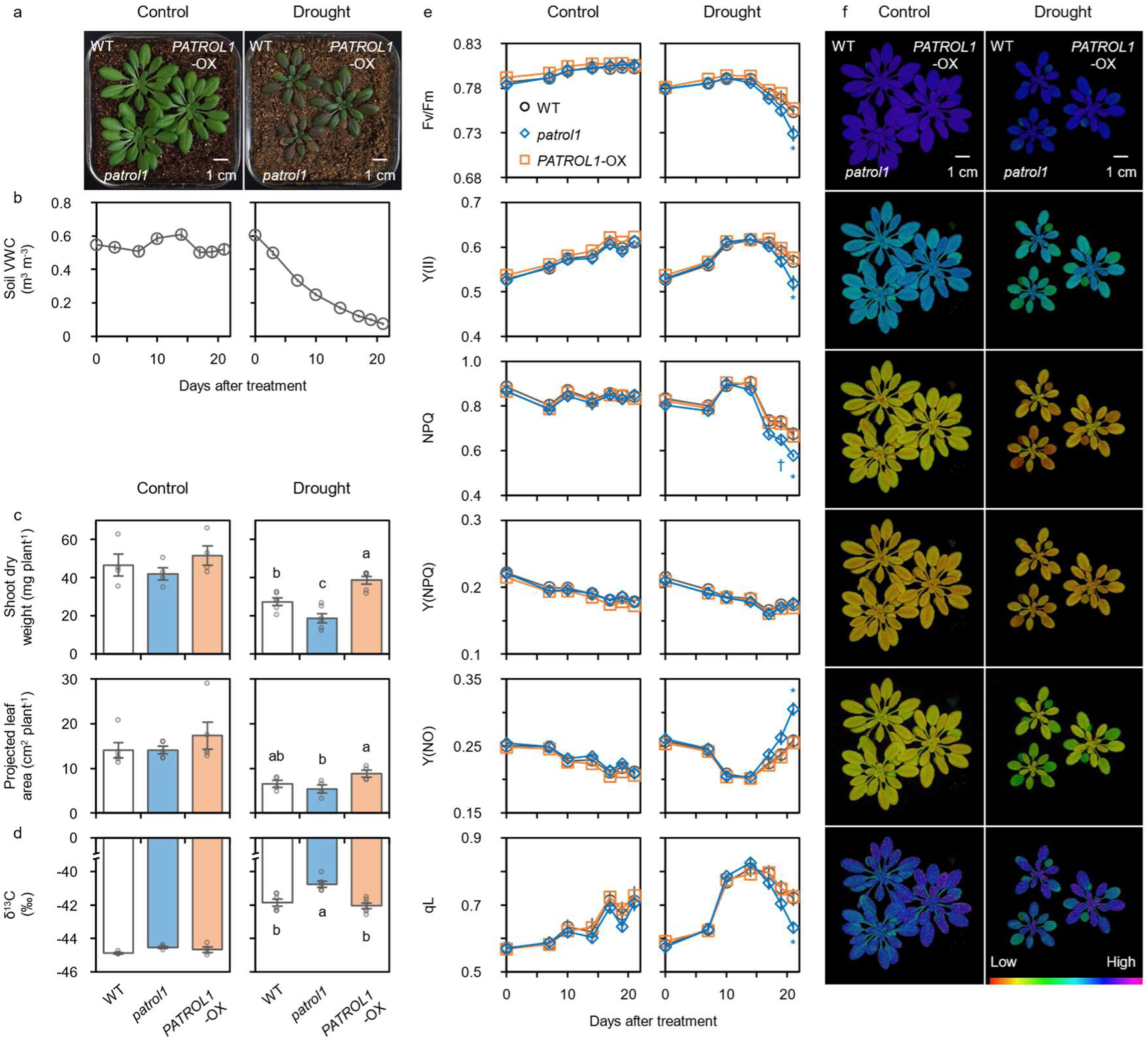
Photochemical activities and growth of *PATROL1* transgenic plants under drought conditions. **a** Phenotype of WT, *patrol1*, and *PATROL1*-OX plants after withholding water for 21 days. **b** Soil volumetric water content (VWC). **c** Shoot dry weight and projected leaf area after treatment. **d** Carbon isotope discrimination (δ^13^C) of harvested shoots. Error bars represent SEM (*n* = 4–6). Means not sharing any letters are significantly different between genotypes at *p* < 0.05 by Tukey–Kramer test. **e** After overnight-dark adaptation, maximum quantum yield of PSII (Fv/Fm), steady-state quantum yield of PSII (Y(II)), non-photochemical quenching (NPQ), quantum yield of regulated thermal dissipation in PSII (Y(NPQ)), that of non-regulated energy dissipation in PSII (Y(NO)), and the fraction of open PSII reaction centers (qL) were measured under PPFD of 100 µmol m^−2^ s^−1^. Error bars represent SEM (*n* = 4–6). Significant differences between WT and transgenic plants at \**p* < 0.05 and †0.10 by Dunnett’s test. **f** Visualization of steady-state photochemical activities under PPFD of 100 µmol m^−2^ s^−1^.

To gain a deeper understanding of the photosynthetic response to soil dehydration, we obtained spatial and time-resolved chlorophyll fluorescence parameters of plants grown at photosynthetically active photon flux density (PPFD) of 100 µmol m^−2^ s^−1^. The maximum quantum yield of PSII (Fv/Fm) and steady-state Y(II) of the treated plants decreased as the drought progressed, especially in *patrol1* (Fig. 4e, f). Concurrently, NPQ of *patrol1* decreased significantly, while the quantum yield of regulated thermal dissipation in PSII (Y(NPQ)) remained comparable among genotypes (Fig. 4e, f). As Y(NPQ) remained stable in *patrol1* plants while Y(II) decreased, excess excitation energy was unrelatedly dissipated as the quantum yield of non-regulated energy dissipation in PSII (Y(NO)) (Fig. 4e), which may have led to the formation of reactive oxygen species ^24,25^. Together, δ^13^C and chlorophyll fluorescence measurements indicate that electron-sink limitation of photosynthesis, induced by stomatal closure from restricted water uptake in the rhizosphere, may have caused *patrol1* to generate reactive oxygen species, leading to photoinhibition (Fig. 4e right), which was more pronounced under higher irradiance (Supplementary Fig. S6).

Finally, to clarify the independent contributions of shoot and root PATROL1 to drought tolerance, we micrografted *A. thaliana* scions and rootstocks with different PATROL1 expression levels. VWC in the control ranged between 0.43 m^3^ m^-3^ and 0.58 m^3^ m^-3^, but under drought decreased from 0.59 m^3^ m^-3^ to 0.04 m^3^ m^-3^ after 24 days. Under drought conditions, plants with *patrol1* as either scion or rootstock (i.e., KO scion/KO rootstock, KO/WT, and WT/KO plants) had significantly lower shoot dry weight, significantly less C content, and smaller leaf area and shoot N content than WT/WT plants (Fig. 5 and Supplementary Fig. S7a). On the other hand, plants with *PATROL1*-OX as either scion or rootstock (i.e., OX/OX, OX/WT, and WT/OX plants) had greater value than WT/WT plants (Fig. 5 and Supplementary Fig. S7a). Consequently, there were strong positive correlations between shoot dry weight, C content, and N content in all genotypes under drought (Fig. 6). In addition, the trend of Y(II) was comparable to shoot growth, being higher in grafted plants including *PATROL1*-OX and lower in plants including *patrol1* (Supplementary Fig. S7b). Under drought, NPQ was lowest in KO/WT, followed by WT/WT, KO/KO, and WT/KO (Supplementary Fig. S7b). Also, the lowest NPQ was observed in WT/OX, followed by OX/OX, OX/WT, and WT/WT. The SPAD value was significantly higher in OX/OX than in WT/WT and lower in WT/KO (Supplementary Fig. S7b). Under control conditions, there were no differences in plant growth or photosynthesis among genotypes (Fig. 5 and Supplementary Fig. S7). In summary, the importance of PATROL1 is comparable between shoots and roots, suggesting that both root uptake and leaf photosynthesis limit plant growth under drought conditions.

**Fig. 5.**
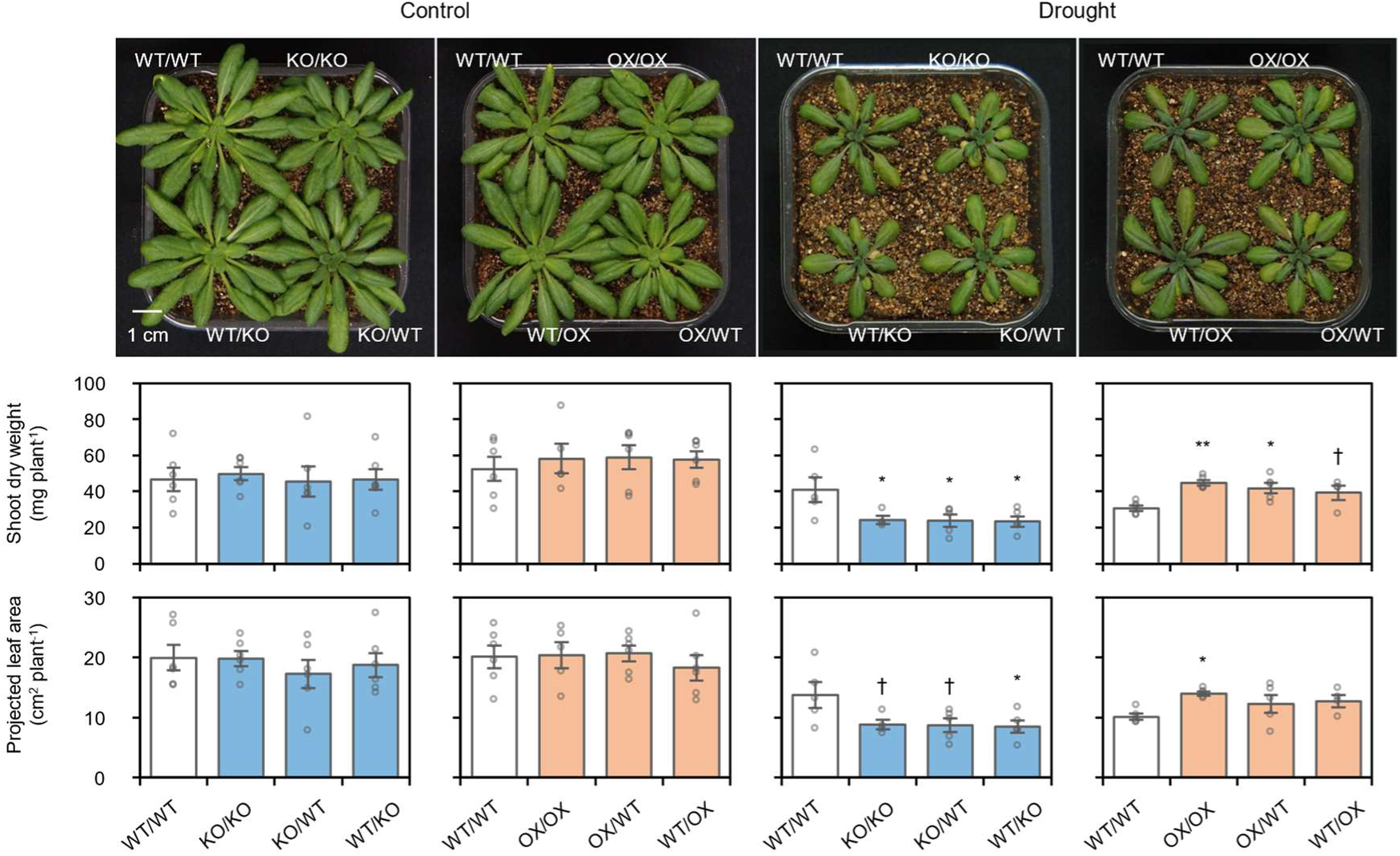
Drought tolerance in grafted plants with different *PATROL1* expression levels in scion and rootstock. Growth performance, shoot dry weight, and projected leaf area of grafted plants after withholding water for 24 days. Error bars represent SEM (*n* = 4–6). Significant differences between WT/WT (scion/rootstock) and transgenic grafted plants at \*\**p* < 0.01, **0.05 and, †0.10 by Dunnett’s test.

**Fig. 6.**
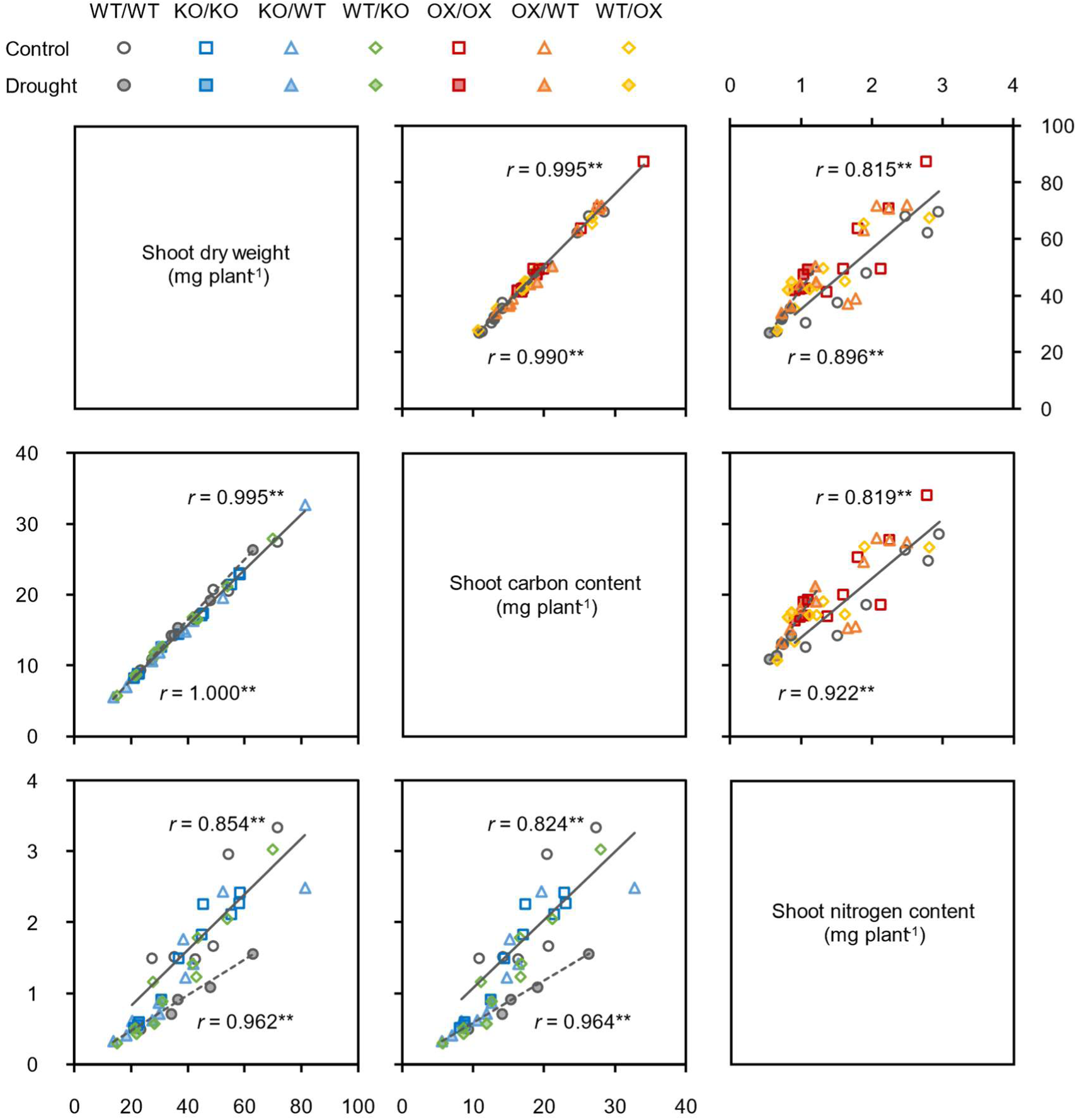
Correlation matrix of shoot dry weight, C content, and N content under drought conditions. ** indicate significant Pearson’s correlation at *p* < 0.01 (*n* = 19– 24).

## Discussion

Enhancing photosynthesis and improving root water and nutrient uptake are essential for achieving drought tolerance and adaptation under anticipated climate change ^2^. Plants may encounter spatial and temporal variations in solar radiation and atmospheric conditions ^4^, soil moisture ^26^, and nutrient availability ^11^. Therefore, it is vital for field-grown crops to have highly plastic root and shoot phenotypes to cope with these challenges ^27^. We showed that overexpression of *PATROL1* enhances photosynthesis and biomass production through vigorous root growth and increased N uptake under water deficit (Fig. 2, 4, 6). We also showed that PATROL1 in roots interacts with AHAs and alters its localization under hyperosmotic stress (Fig. 1, 3). These results indicate that PATROL1 regulates PM H^+^-ATPase localization in response to fluctuating environmental conditions (e.g., water and nutrients availability), not only in leaves ^18,23,28^ but also in roots. We previously demonstrated that rapid stomatal movement of *PATROL1*-OX plants improved photosynthesis and biomass gain under fluctuating light and adequate irrigation ^21^. Thus, optimizing PM H^+^-ATPase trafficking in both shoots and roots by modulating the single gene *PATROL1*, could serve as an adaptive strategy to improve plant growth in both optimal and sub-optimal environments.

The present GUS reporter assay confirmed that *PATROL1* is ubiquitously expressed throughout the plant (Fig. 1b), which collaborates with previous studies ^18,22^. The confocal microscopic analysis confirmed that the PATROL1 level is notably high in the meristematic regions of shoots and primary roots, lateral root primordia, and root stelae (Fig. 1c), where it plays a key role in root growth and water and nutrient foraging ^11^. In addition, as the maximum CO_2_ assimilation rate and Y(II) of *patrol1* plants remained suppressed even under high [CO_2_] (Supplementary Fig. S2), indicating that reduced stomal conductance is not the sole cause of the growth reduction (Fig. 1a). Overall, these results raise the question of how PATROL1 in both shoots and roots contributes to plant growth. Micrografting between WT and *patrol1* or *PATROL1*-OX plants revealed that PATROL1 is indispensable in both shoots and roots, indicating that root uptake and leaf photosynthesis are simultaneous limiting factors for plant growth under drought conditions (Fig. 5). Grafted plants with higher transpiration demand than their root water supply (WT/KO, OX/WT) had greater NPQ and less linear electron flow activity than combinations with the converse (KO/WT, WT/OX) under drought (Supplementary Fig. S7b). This indicates that effective root water and nutrient foraging are essential for preventing dehydration and maintaining photosynthesis during soil drying. However, the reduced shoot growth in KO/WT plants and the lack of growth enhancement in WT/OX relative to WT/WT suggest that carbon gain from photosynthesis is necessary for shoot growth alongside root improvement (Fig. 5), as root growth also depends on sufficient C allocation ^4^. Overall, *PATROL1* overexpression in both roots and shoots (OX/OX) enhanced shoot growth, N and C assimilation, and chlorophyll content (Fig. 5 and Supplementary Fig. S7), emphasizing the need for simultaneous enhancement of root functions and leaf photosynthesis to improve biomass production under drought. Ultimately, *PATROL1* overexpression throughout the plant enhanced drought tolerance, resulting in increased photosynthesis and shoot biomass relative to WT under moderate drought conditions (Fig. 4 and Supplementary Fig. S6).

Coimmunoprecipitation of AHAs with PATROL1 (Fig. 1d) suggests a direct or indirect interaction between them in roots. PATROL1 may contribute to drought tolerance by facilitating membrane trafficking, particularly the delivery of PM H^+^-ATPase ^18^. PATROL1 is related to mammalian uncoordinated 13s (Munc13s), which are involved in tethering, docking, and rapid fusion of transport vesicles and target membranes through interactions with SNAREs (soluble *N*-ethylmaleimide-sensitive factor attachment protein receptors) ^29^. PATROL1 contains a MUN domain (Supplementary Fig. S1), a critical region also found in Munc13-1 ^30,31^. Recent studies have revealed that SNAREs also interact with AHAs ^32–36^, and potentially with PATROL1, as indicated by immunoprecipitation mass spectrometric analysis ^37^. In this context, PATROL1 could ensure proper trafficking of PM H^+^-ATPase either via SNAREs or directly. Moreover, the altered localization of PATROL1-GFP bodies in the root epidermal cells in response to PEG treatment (Fig. 3b), is consistent with previous studies ^18,22,23,28^, as well as transcriptional regulation (Fig. 3a). Together with the co-IP result (Fig. 1b), these results support the hypothesis that PATROL1 in roots colocalizes with and possibly regulates PM H^+^-ATPase activity in response to hyperosmotic stress.

There is a growing recognition of the roles of PM H^+^-ATPase in root system plasticity and nutrient uptake under various conditions ^12^. Here, we show that under PEG-induced water deficit, *PATROL1*-OX plants had significantly greater total root length and lateral root numbers (Fig. 2b, c) than WT and *patrol1* plants. Furthermore, N accumulation tended to decrease in *patrol1* and increase in *PATROL1*-OX plants under drought (Supplementary Fig. S7a), resulting in significant strong positive correlations between shoot N, C, and growth (Fig. 6). Although we did not confirm the colocalization of AHAs, the distribution patterns of PATROL1 in the primary root tip, mature zone, and lateral root tip (Fig. 1c and Supplementary Fig. S3) closely resemble those of AHA1, AHA2, and AHA7 ^38–41^. It was reported that greater H^+^ secretion by PM H^+^-ATPase in the root tip sustains root elongation and root hair growth under moderate water stress in both *A. thaliana* and *O. sativa* ^42^. A recent study using bromocresol purple assessment showed that *patrol1* roots released less H^+^ than WT roots, while *PATROL1*-OX roots released more ^22^. However, the similarity of expression patterns of *AHA* family members in WT, *patrol1*, and *PATROL1*-OX across all lines (Supplementary Fig. S5) suggests that PATROL1 regulates PM H^+^-ATPase activity post-transcriptionally rather than transcriptionally. Downregulation of AHA1, AHA2, and AHA7 leads to auxin hypersensitivity and inhibits root elongation, while constitutive AHA1 activation has the opposite effect ^43^. Live imaging shows AHA2 localized at the root surface, with stronger signals at the root tip, and its accumulation in the cytoplasm reduces H^+^ secretion when root elongation is suppressed ^38^. AHA2 also promotes root branching and growth in response to N availability ^39,44^, and enhances hydrotropism under hyperosmotic stress ^45,46^. Constitutive AHA1 activation enhances root elongation, increases shoot mineral contents (e.g., N, K, Ca, S, P), and boosts shoot biomass under nutrient-deficient conditions ^47^. Similarly, overexpression of *O. sativa* H^+^-ATPase 1 in rice enhances root nutrient uptake and transportation, increasing leaf N, K, P, and Ca ^48^. Recent research also indicates that PM H^+^-ATPase activity contributes to nitrate assimilation in *A. thaliana* leaves ^49^. Collectively, these findings suggest that PM H^+^-ATPase plays a crucial role in nutrient uptake and transport by proton motive force ^13^. The consistency of enhanced root growth, increased shoot N content, and improved drought tolerance in *PATROL1*-OX plants with known functions of PM H^+^- ATPase indicates that PATROL1 promotes root plasticity and activity under soil water stress by regulating PM H^+^-ATPase activity. In addition, since cellulose synthase complexes are another identified cargo of PATROL1-mediated plasma membrane trafficking ^50^, root elongation could also be linked to cellulose biosynthesis.

In conclusion, we have uncovered a new function of the membrane trafficking factor PATROL1 in root system plasticity, N uptake, and biomass production in response to water deficit. As PATROL1 is conserved among higher plants ^18^, our findings suggest that enhancing PM H^+^-ATPase regulation through PATROL1 overexpression could be a promising strategy for developing drought-tolerant crops. Further research on the interactions between PATROL1, PM H^+^-ATPase, and candidate proteins such as SNAREs will be crucial to elucidating the mechanisms underlying these phenotypes.

## Method

### Plant materials and growth conditions

We used *Arabidopsis thaliana* wild type (Col-0), a *patrol1* knockout mutant, and *p35S:GFP-PATROL1; patrol1* overexpression lines (*PATROL1*-OX) ^18^. We generated *gPATROL1:GUS-PATROL1; patrol1* (*gPATROL1:GUS-PATROL1*) and *gPATROL1:EGFP-PATROL1; patrol1* (*gPATROL1:EGFP-PATROL1*) transgenic lines. Sterilized seeds were planted in 380-mL pots filled with a mixture of equal volumes of vermiculite and nutrient soil (Metromix 350; Sun Gro Horticulture, USA). The plants were grown in a growth chamber (NK system, Japan) at 23°C with a relative humidity (RH) of 65%, a photosynthetically active photon flux density (PPFD) of 100 µmol m^−2^ s^−1^, and an atmospheric CO_2_ concentration ([CO_2_]) of 400 µmol mol^−1^ under a 10-h light period. For elevated [CO_2_] experiment, plants were grown at a [CO_2_] of 1500 µmol mol^−1^ under an 8-h light period.

Drought stress was induced by withholding water for 21 days from 16-day-old intact plants grown in soil and for 24 days from 27-day-old grafted plants. Soil volumetric water content (VWC) was determined every 2 to 4 days by the gravimetric method, where:

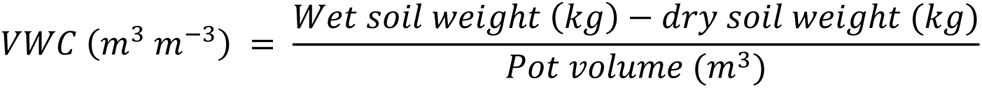

For hyperosmotic stress treatment, 5-day-old seedlings were transferred to vertically oriented solid medium (1/2 MS, 1% sucrose, 0.05% MES, pH 5.7, 0.8% gellan gum), which was first dehydrated with 30% (w/v) PEG 6000 (FUJIFILM Wako Pure Chemical, Japan) ^51^, and grown for an additional 10 days.

### Micrografting

Transgenic plants were micrografted as described ^52,53^, with modifications. In brief, seeds were surface-sterilized, soaked in distilled water, and kept in the dark at 4°C for 3 days to synchronize germination. They were sown on a square petri dish containing 1% agar medium (1/2 MS, 1% sucrose, 0.05% MES, pH 5.7) and grown vertically in the dark at 23°C for 2 days. The petri dishes were then moved to a 10-h light period with a PPFD of 100 µmol m^−2^ s^−1^ for 2 days. Next, hypocotyls were cut with a disposable razor blade (FA-10; FEATHER Safety Razor, Japan) and 90°-butt-grafted ^52^ on a nylon membrane (RPN119B; GE Healthcare, USA) placed on top of 2% agar medium (1/2 MS, 1% sucrose, 0.05% MES, pH 5.7). The grafted plants were incubated at a 45° angle under a PPFD of 50 µmol m^−2^ s^−1^ for 2 days, then 100 µmol m^−2^ s^−1^ for 4 days. Adventitious roots from the scion were removed, and the grafted plants were transferred to 1% agar medium (without nylon membrane) for an additional 5 days. Finally, adventitious roots were removed again before the grafted plants were transplanted into pots filled with soil.

### Plasmid construction

To prepare the construct comprising PATROL1 fused to GUS or EGFP at the N-terminus (GUS-PATROL1 or EGFP-PATROL1) under the control of the *PATROL1* regulatory sequence, we amplified the 5’-upstream region of *PATROL1* and the coding sequence of *GUS* or *EGFP* by PCR using appropriate templates and cloned together into the PstⅠ/EcoRⅠ-digested pRI201AN vector (Takara Bio, Japan). The resulting plasmids were digested with SmaⅠ, and PCR-amplified genomic fragments containing the coding region and the 3′-region of *PATROL1* were inserted into the SmaⅠ-digested vectors, to construct *gPATROL1:GUS-PATROL1* or *gPATROL1:EGFP-PATROL1*. Primers used for plasmid construction are listed in Supplementary Table 1.

### Transformation of *A. thaliana*

*Agrobacterium tumefaciens* strain GV3101 was transformed with the appropriate construct and then used to transform *A. thaliana patrol1* mutants by the floral-dip method ^54^.

### Co-immunoprecipitation analysis

Approximately 300 mg of 7-day-old *A. thaliana* roots expressing either GFP-PATROL1 ^18^ or GFP alone was ground in liquid nitrogen and homogenized in 900 μL of cold lysis buffer (50 mM Tris-HCl, pH7.5, 150 mM NaCl, 1 mM EDTA, 1% Triton X-100, and a protease inhibitor cocktail). The homogenates were centrifuged at 15,000×g for 10 min at 4°C. The supernatants were immunoprecipitated with a μMACS GFP Isolation Kit (Miltenyi Biotec, Japan), according to the manufacturer’s instructions. The immunoprecipitates were separated by SDS-PAGE and immunoblotted with an anti-GFP antibody (sc-9996, 1:1,000 dilution; Santa Cruz Biotechnology, USA) or an anti-AHA2-cat polyclonal antibody (1:3000 dilution; kindly provided by Dr. Toshinori Kinoshita ^55^). Secondary antibodies—anti-Mouse IgG, HRP-Linked Whole Ab Sheep (NA931, 1:10000 dilution; GE Healthcare, USA), and Anti-Rabbit IgG, HRP-Linked Whole Ab Donkey (NA934, 1:10000 dilution; GE Healthcare, USA)—were used to detect the primary antibodies on SuperSignal West Dura Extended Duration Substrate (34076; Thermo Fisher Scientific, USA). A protein–protein BLAST search showed that the AHA2 antibody recognition sequence ^55^ shared high identity with those of all AHA isoforms (Supplementary Table 2).

### Histochemical analysis of GUS staining

Two-week-old *gPATROL1:GUS-PATROL1* plants were pre-fixed in ice-cold 90% (v/v) acetone for 15 min. The fixed samples were rinsed several times in sterile deionized water and incubated in staining buffer (100 mM sodium phosphate buffer (pH 7.0), 0.1% Triton-X-100, 5 mM potassium ferrocyanide, 5 mM potassium ferricyanide, 10 mM EDTA, 0.5 mg/mL 5-bromo-4-chloro-3-indolyl-β-D-glucuronic acid) for 12 h at 37°C. Stained samples were rinsed several times in 70% ethanol and transferred to chloral hydrate clearing solution (chloral hydrate:water:glycerol = 8:1:3 [w/v/v]) for observation.

### Quantitative reverse-transcription PCR (RT-qPCR)

Short-term effects of osmotic stress were observed by pouring 50 mL 30% (w/v) PEG 6000 onto 19-day-old *A. thaliana* grown on 1/2 MS gellan gum medium. Total RNA from shoots and roots was extracted with an RNeasy Plant Mini Kit (QIAGEN, Germany) and treated with DNase (TURBO DNA-free Kit; Thermo Fisher Scientific, USA), and cDNA was synthesized using the High Capacity RNA-to-cDNA Kit (Thermo Fisher Scientific, USA). Power SYBR Green PCR Master Mix (Thermo Fisher Scientific, USA) was used for RT-qPCR, and the 2^−ΔΔCT^ method was used to calculate expression levels relative to *ACTIN 2* as an internal standard. Relative expression levels of all *A. thaliana* Domain of Unknown Function 810 (DUF810) and AHA member genes in the shoots and roots of 14-day-old plants grown on 1/2 MS gellan gum medium were similarly determined. Sequences of primers used for RT-qPCR are listed in Supplementary Table 1.

### Confocal microscopic analysis

Confocal microscopic analysis of EGFP-PATROL1 was performed using a C2 confocal Microscope (Nikon, Japan). With excitation at 488 nm, EGFP fluorescence was measured at 500-550 nm. Images were processed in NIS-elements (Nikon, Japan) and ImageJ/Fiji software ^56^.

Roots of 1-week-old *gPATROL1:EGFP-PATROL1* transgenic plants, grown on the vertically oriented 1/2 MS gellan gum medium, were mounted between two different-sized coverslips 24 × 60 mm and 24 × 45 mm, with a sheet of Parafilm as a spacer, and the gap was filled with liquid 1/2 MS medium (sucrose-free, pH 5.7). For osmotic-stress treatments, seedlings were perfused with liquid 1/2 MS medium with or without 10% (w/v) PEG 8000 (P5413; Sigma-Aldrich, USA) and incubated under the microscope for 30 min. Root epidermal cells were imaged through an inverted confocal microscope (FV1000; Olympus, Japan) with an air objective UPLSAPO 40X (NA 0.95, WD 0.18). GFP was excited at 473 nm with a multi-line argon laser. Confocal Z-stacked images were acquired at 1-µm intervals before and after treatment. Images were processed and analyzed in ImageJ/Fiji software.

### Root growth assay

Photographs were taken 10 days after PEG treatment with a digital single-lens reflex camera. Root length and numbers were manually measured in the ImageJ/Fiji software ^56^.

### Gas exchange and chlorophyll fluorescence measurements

We measured gas exchange and chlorophyll fluorescence of WT, *patrol1*, and *PATROL1*-OX plants grown under elevated [CO_2_] (1500 μmol mol^−1^) using a portable gas exchange system (LI-6400XT; LI-COR Biosciences, USA). Following overnight-dark adaptation, we measured the light response of the CO_2_ assimilation rate (*A*), the quantum yield of PSII (Y(II)), and non-photochemical quenching (NPQ) at ambient and elevated [CO_2_] (400 and 1500 μmol mol^−1^), and measured post-illumination induction of *A*, Y(II), NPQ, stomatal conductance (*g_s_*), intercellular CO_2_ concentration (*C_i_*), and intrinsic water use efficiency (WUE*_i_*) at elevated [CO_2_] under PPFD of 500 µmol m^−2^ s^−1^.

Following overnight dark adaptation, chlorophyll fluorescence was determined by a pulse-amplitude-modulated fluorometer (IMAG-MAX/L; Heinz Walz, Germany) under growing light (GL, PPFD 100 µmol m^−2^ s^−1^) and high light (HL, PPFD 500 µmol m^−2^ s^−1^) intensities. Fluorescence was measured every 30 s after light irradiation, and we calculated the maximum quantum yield of PSII (Fv/Fm), Y(II), NPQ, quantum yield of regulated thermal dissipation in PSII (Y(NPQ)), that of non-regulated energy dissipation in PSII (Y(NO)), and fraction of open PSII reaction centers (qL) as described ^24,57^.

### Measurements of Carbon isotope discrimination (δ^13^C)

δ^13^C of harvested shoots was determined with a CN analyzer (Vario Micro; Elementar Analyzensysteme, Germany) connected to an isotopic ratio mass spectrometer (IsoPrime 100; Isoprime, UK). δ^13^C values (‰) were calculated as described ^58^.

### Determination of total carbon and nitrogen contents

Harvested shoots oven-dried at 80°C for 72 h to a constant weight and weighted. Samples were ground to a fine powder, and total C and N contents were determined by element analyzer (SUMIGRAPH NC-22F; Sumika Chemical Analysis Service, Japan).

### Statistical Analysis

All *n* values and statistical tests used are indicated in the figure legends. Data were analyzed by two-tailed *t*-test, or by one-way ANOVA with Dunnett’s or Tukey’s *post-hoc* test at *p* < 0.05 in R v. 4.3.1 software (https://www.r-project.org/).

## Author contributions

N.K., K.S., H.K., and W.Y. conceived and designed the study. N.K., K.S., H.K., Y.Shimizu, Y.S., K.N., and M.A. conducted experiments and performed data analysis. I.T and W.Y. contributed to the original idea of the project and supervised the study. N.K., Y.Shimizu, Y.S., K.N., M.A., I.T., and W.Y. wrote the manuscript. All authors discussed the results and revised the manuscript.

## Data availability

Supporting data can be requested by contacting the corresponding author.

## Acknowledgement

We thank Dr. Hashimoto-Sugimoto Mimi and Dr. Iba Koh (Kyusyu University) for providing the seeds, Dr. Toshinori Kinoshita (Nagoya University) for supplying the AHA2 antibody, Dr. Seo Mitsunori (University of the Ryukyus) for guidance on micrografting, and Dr. Taneda Haruhiko, Dr. Masaru Kono, and Dr. Qu Yuchen (University of Tokyo) for their technical support with carbon isotope discrimination measurements and for their valuable discussions.

---

**Supplementary Fig. S1.**
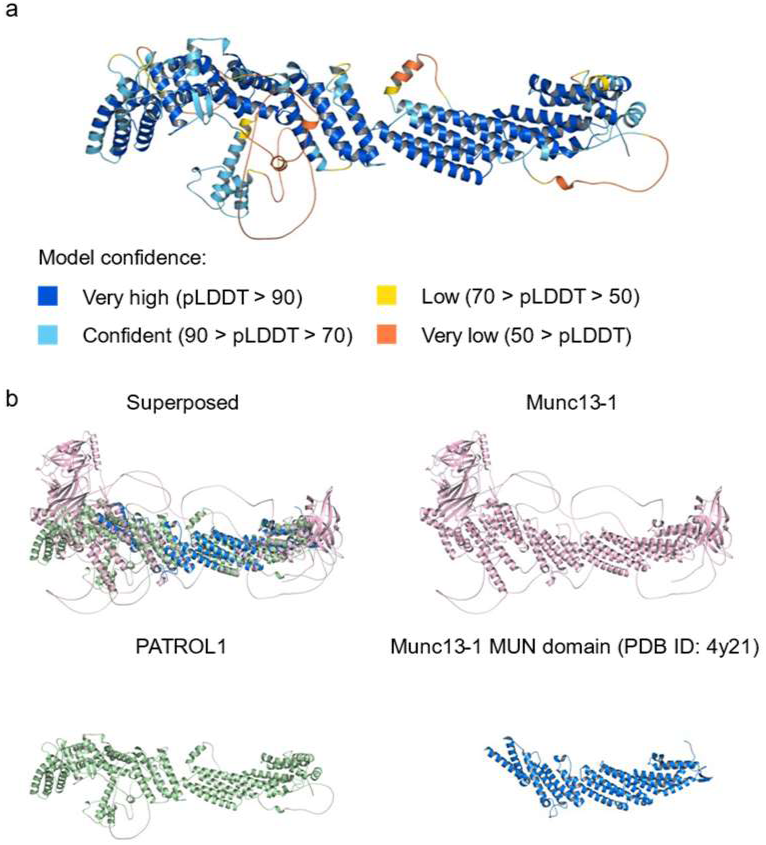
Comparison of PATROL1 and Munc13 protein structures. **a** Model of predicted *A. thaliana* PATROL1 structure from AlphaFold Protein Structure Database. Each residue of the protein is color-coded by its corresponding per-residue model confidence score (pLDDT). **b** Superposition of PATROL1 (AlphaFold) with the *Homo sapiens* Munc13-1 (AlphaFold) and *Rattus norvegicus* Munc13-1 C1-C2B-MUN-C2 C domain (PDB ID: 7t7v) by the PyMOL Molecular Graphics System, Version 2.5.0 (Schrodinger, USA).

**Supplementary Fig. S2.**
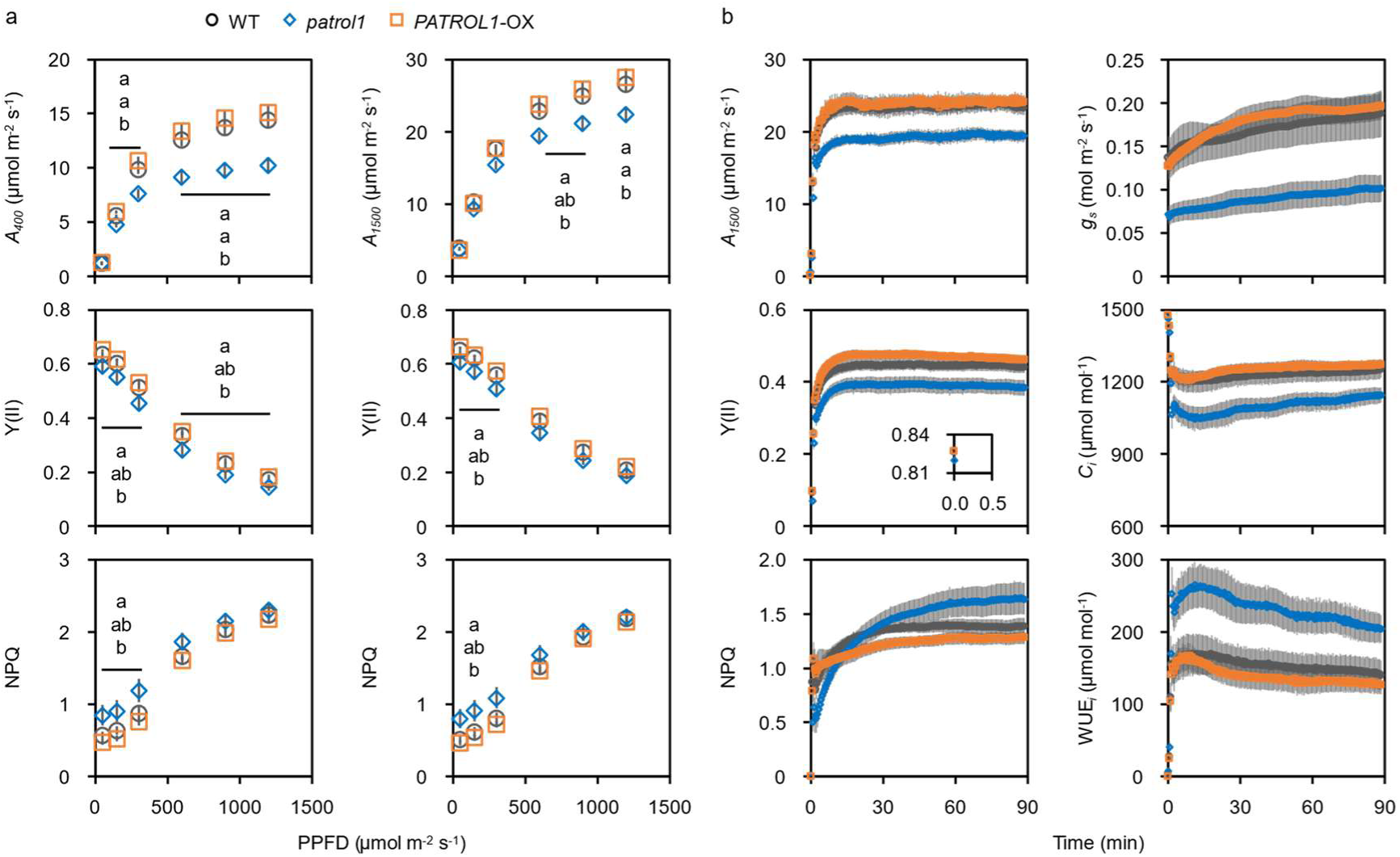
Photosynthetic properties of *PATROL1* transgenic plants grown under elevated [CO_2_]. **a** Light response of CO_2_ assimilation rate (*A*), quantum yield of PSII (Y(II)), and non-photochemical quenching (NPQ) at ambient and elevated [CO_2_] (400 and 1500 μmol mol^−1^). Error bars represent SEM (*n* = 6–7). Means not sharing any letters are significantly different between genotypes at *p* < 0.05 by Tukey– Kramer test. **b** After overnight dark adaptation, post-illumination induction of *A*, Y(II), NPQ, stomatal conductance (*g_s_*), intercellular CO_2_ concentration (*C_i_*), and intrinsic water use efficiency (WUE*_i_*) at elevated [CO_2_] (1500 μmol mol^−1^) measured under PPFD of 500 µmol m^−2^ s^−1^. Error bars represent SEM (*n* = 6–7).

**Supplementary Fig. S3.**
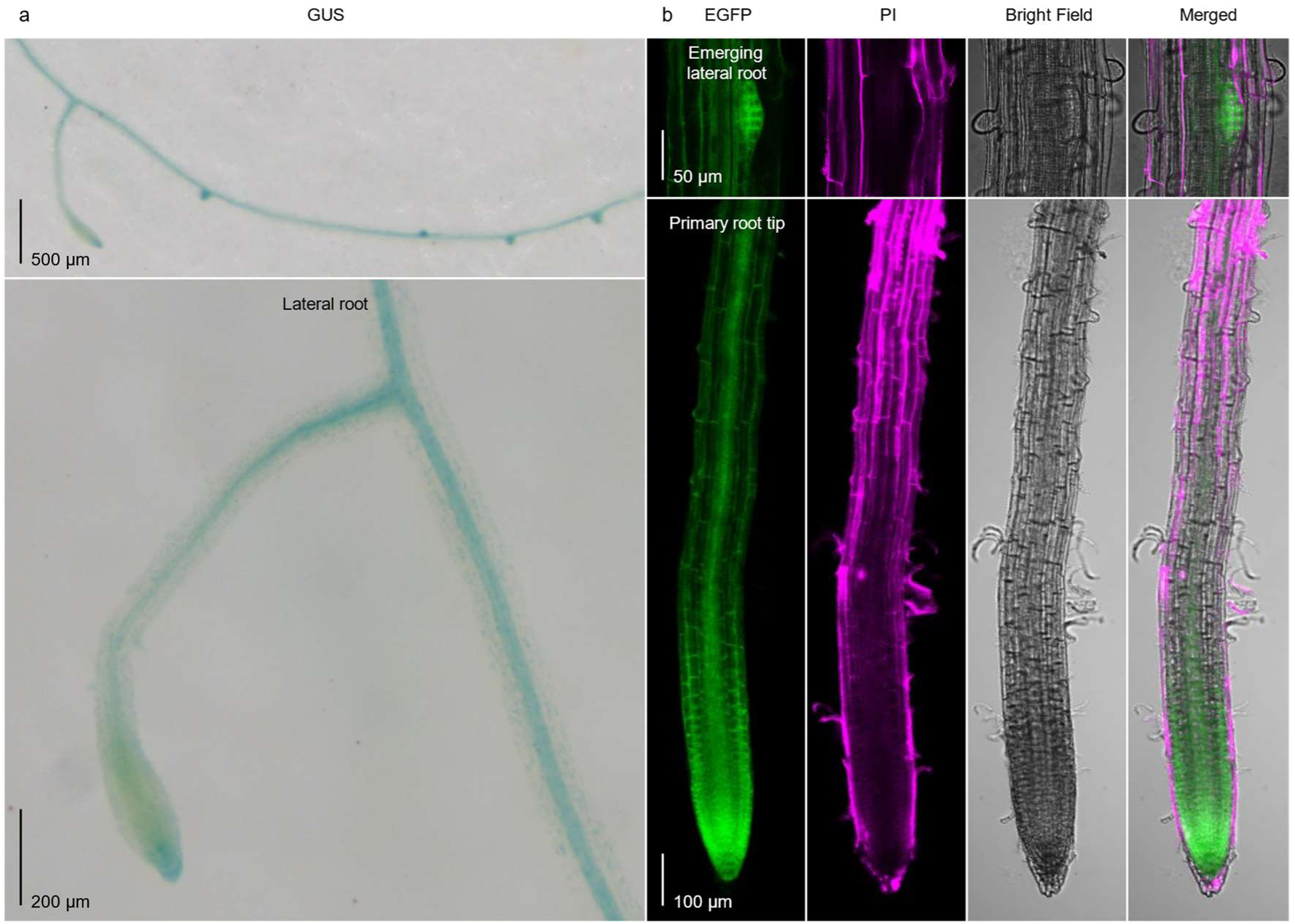
Localization of PATROL1 in roots of *A. thaliana*. **a** GUS staining of 2-week-old *A. thaliana* root producing GUS-PATROL1. **b** Localization of EGFP-PATROL1 in 2-week-old primary root. Green, EGFP-PATROL1 localization; magenta, cell wall propidium iodide (PI) staining.

**Supplementary Fig. S4.**
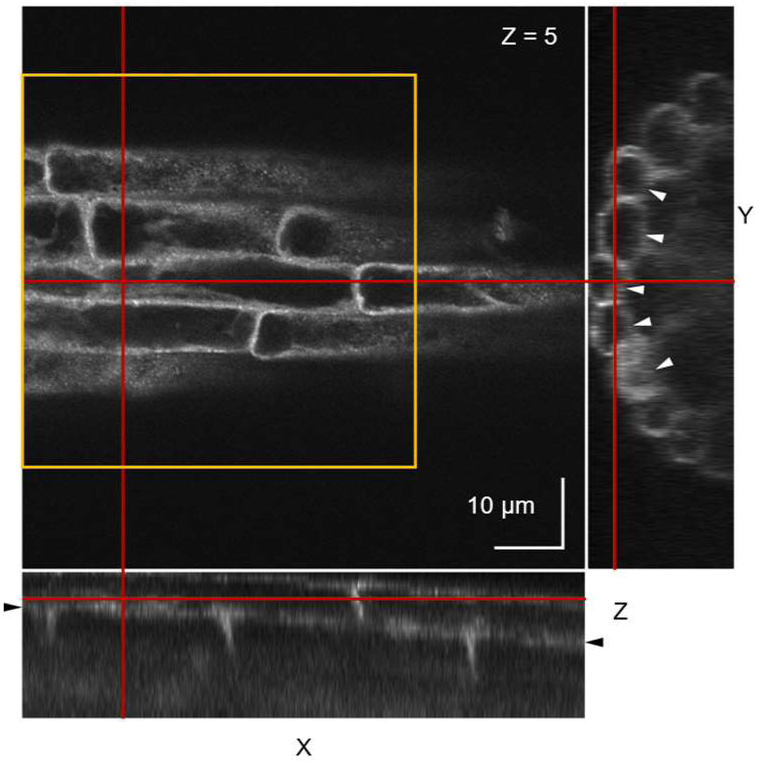
Representative 3D reconstruction (Z-stack) image of EGFP-PATROL1 in root epidermal cells of *A. thaliana* after PEG treatment. Orthogonal views at Z = 5 were generated on a Z-stacked image of 23 slices with 1-µm intervals. The X-Y and Y-Z views show that EGFP-PATROL1 is localized in the cytosol along the plasma membrane of root epidermal cells. The X-Z and Y-Z views indicate that slices Z = 1–5 contain the centrifugal-side signal, but not the centripetal-side signal (arrowheads). The maximum projection image in Fig. 3b was generated using slices Z = 1–5 in the yellow square. Red lines = orthogonal axes. Bars = 10 mm.

**Supplementary Fig. S5.**
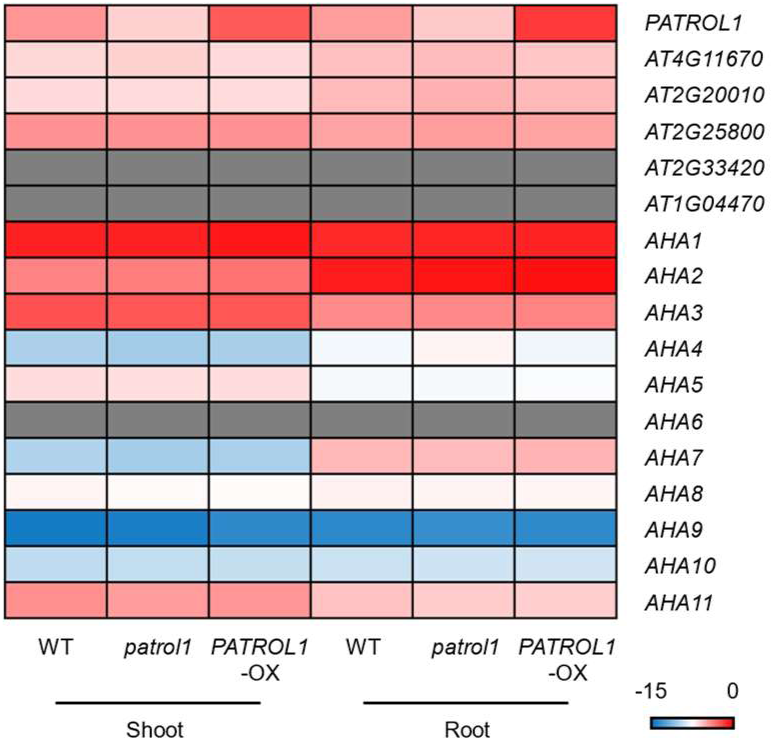
Tissue-specific expression patterns of *A. thaliana* DUF810 and AHA member genes in *PATROL1* transgenic plants. The expression levels of all genes were determined by RT-qPCR and normalized to *ACTIN 2* as an internal standard, and log2 ratio were calculated (*n* = 4). Gray rows represent undetected expression levels.

**Supplementary Fig. S6.**
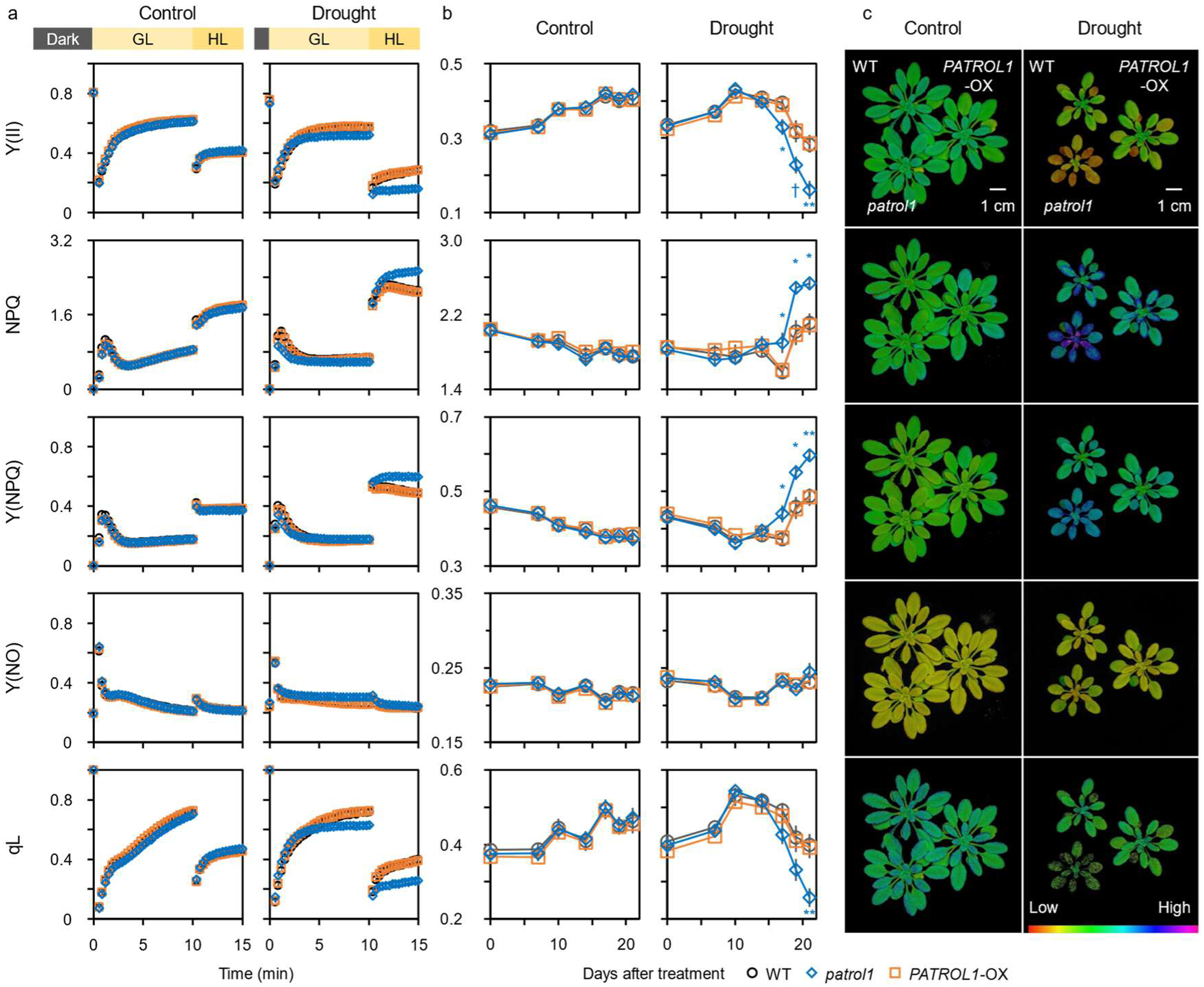
Photochemical activities of *PATROL1* transgenic plants after 21 days of drought treatment. **a** After overnight dark adaptation, post-illumination induction of the quantum yield of PSII (Y(II)), non-photochemical quenching (NPQ), quantum yield of regulated thermal dissipation in PSII (Y(NPQ)), that of non-regulated energy dissipation in PSII (Y(NO)), and the fraction of open PSII reaction centers (qL) were measured under growing light (GL, PPFD of 100 µmol m^−2^ s^−1^) and high light (HL, PPFD of 500 µmol m^−2^ s^−1^) intensities. Error bars represent SEM (*n* = 4–6). **b** Significant differences between WT and transgenic plants at \*\**p* < 0.01, *0.05, and †0.10 by Dunnett’s test. **c** Visualization of steady-state photochemical activities under PPFD of 500 µmol m^−2^ s^−1^.

**Supplementary Fig. S7.**
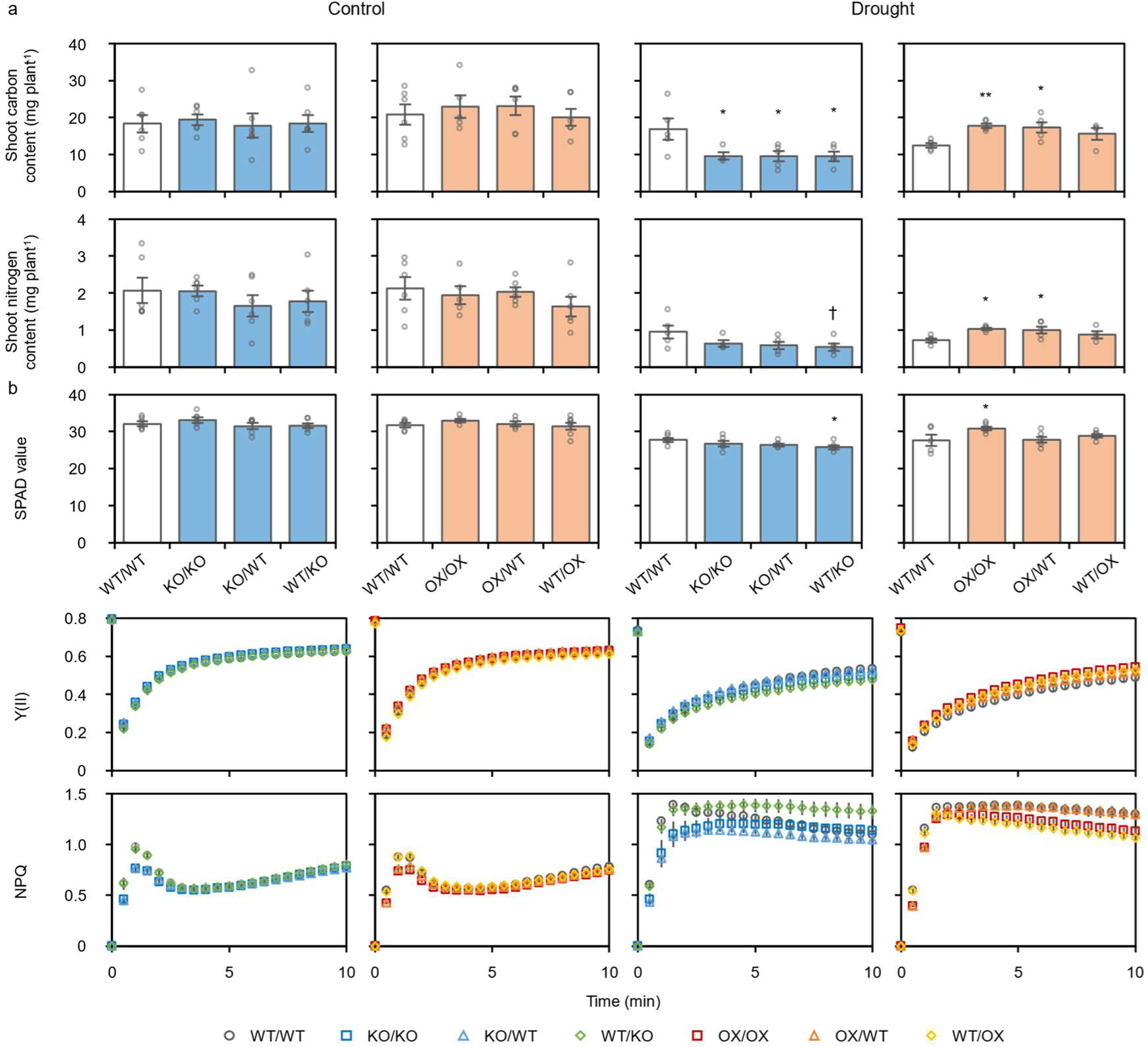
Shoot C, N content, and photochemical activities of grafted plants with different *PATROL1* expression levels in scion and rootstock. **a** Shoot C and N contents of grafted plants. **b** SPAD value; after 14 h of overnight-dark adaptation, post-illumination induction of the quantum yield of PSII (Y(II)) and non-photochemical quenching (NPQ) were measured under PPFD of 100 µmol m^−2^ s^−1^. Error bars represent SEM (*n* = 4–6). Significant differences between WT/WT (scion/rootstock) and transgenic grafted plants at \*\**p* < 0.01, *0.05 and, †0.10 by Dunnett’s test.

**Supplementary Table 1.**
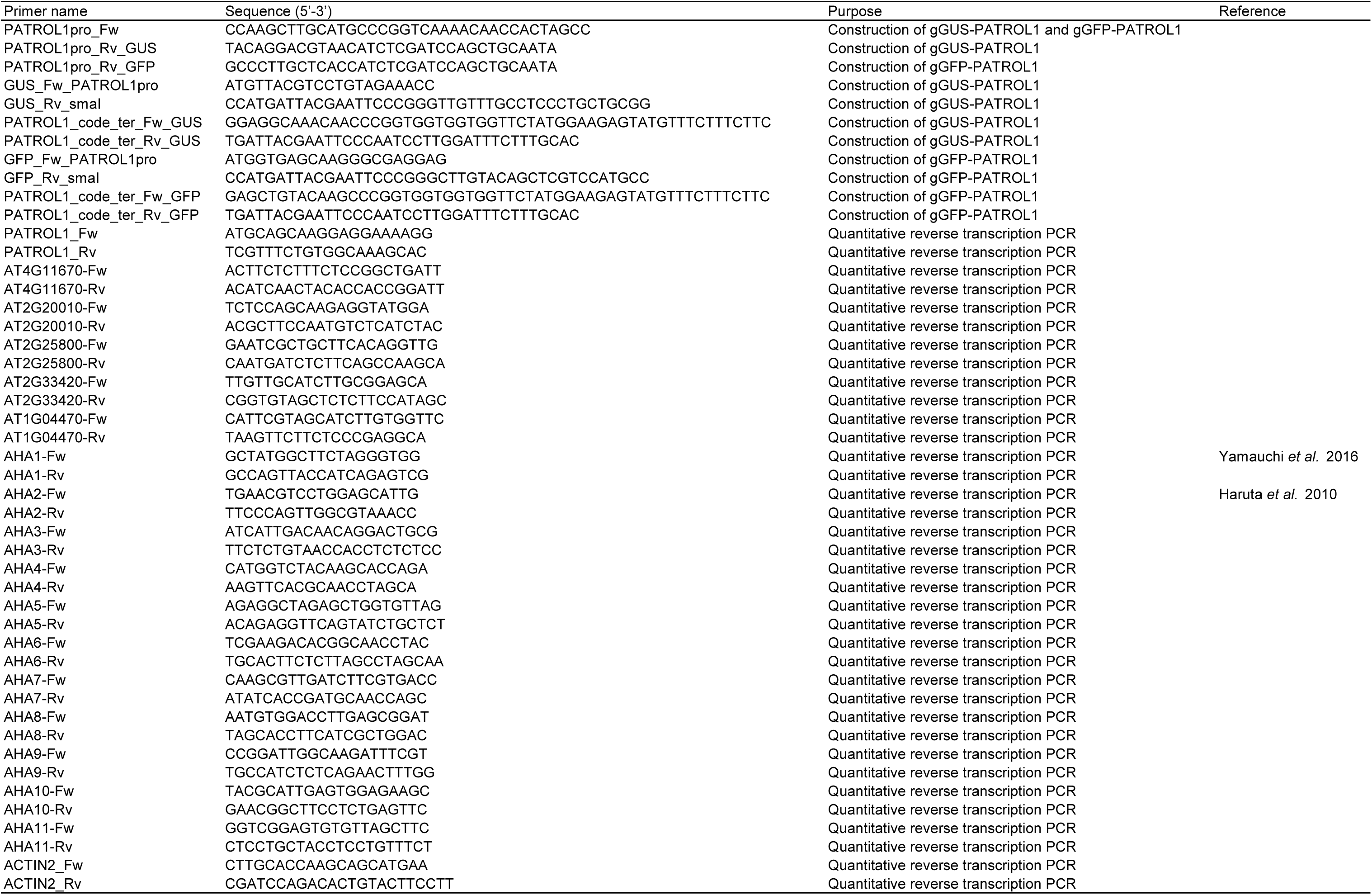
List of primers used in this study.

**Supplementary Table 2.**
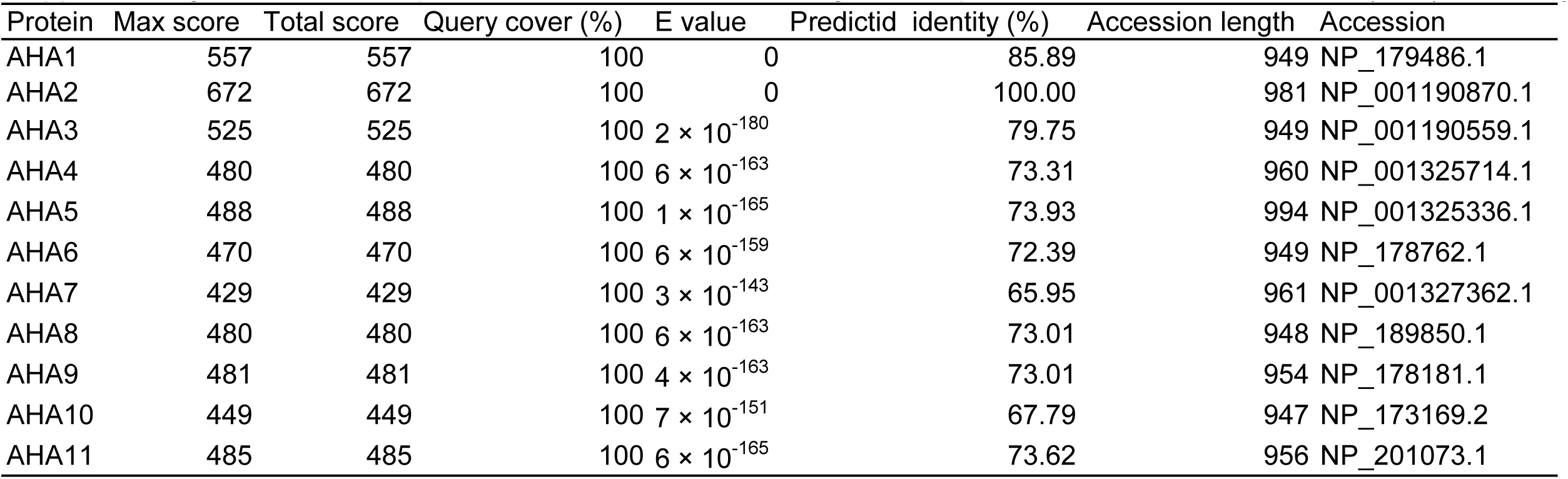
Protein–protein BLAST results for the recognition sequence of *A. thaliana* H^+^-ATPase (AHA) 2 antibody across all AHA members.

